# Neural Correlates of Positive and Negative Valence System Dysfunction in Adolescents Revealed by Data-Driven Parcellation and Resting-State Network Modeling

**DOI:** 10.1101/2020.03.20.001032

**Authors:** Vilma Gabbay, Qi Liu, Samuel J. DeWitt, Lushna M. Mehra, Carmen M. Alonso, Benjamin A. Ely

**Affiliations:** Dept. of Psychiatry & Behavioral Sciences, Albert Einstein College of Medicine, The Bronx, NY; Nathan S. Kline Institute for Psychiatric Research, Orangeburg, NY; Dept. of Psychology, Florida State University, Tallahassee, FL

**Author notes:** **Corresponding Author:** Name: Vilma Gabbay, MD, Address: Van Etten 4A-44, 1225 Morris Park Ave, The Bronx, NY.

**Keywords:** Graph Theory, RDoC, Depression, Anxiety, Anhedonia, fMRI

## Abstract

**Objective:** Adolescence is a period of rapid brain development when symptoms of mood, anxiety, and other disorders often first emerge, suggesting disruptions in maturing reward circuitry may play a role in mental illness onset. Here, we characterized associations between resting-state network properties and psychiatric symptomatology in medication-free adolescents with a wide range of symptom severity.

**Methods:** Adolescents (age 12-20) with mood and/or anxiety symptoms (n=68) and healthy controls (n=19) completed diagnostic interviews, depression/anhedonia/anxiety questionnaires, and 3T resting-state fMRI (10min/2.3mm/TR=1s). Data were preprocessed (HCP Pipelines), aligned (MSMAll), and parcellated into 750 nodes encompassing the entire cortex/subcortex (Cole-Anticevic Brain-wide Network Partition). Weighted graph theoretical metrics (Strength Centrality=C_Str_; Eigenvector Centrality=C_Eig_; Local Efficiency=E_Loc_) were estimated within *Whole Brain* and task-derived *Reward Anticipation*/*Attainment*/*Prediction Error* networks. Associations with clinical status and symptoms were assessed non-parametrically (two-tailed *p_FWE_*<0.05).

**Results:** Relative to controls, clinical adolescents had increased ventral striatum C_Eig_ within the *Reward Attainment* network. Across subjects, depression correlated with subgenual cingulate C_Str_ and E_Loc_, anhedonia correlated with ventromedial prefrontal C_Str_ and lateral amygdala E_Loc_, and anxiety negatively correlated with parietal operculum C_Eig_ and medial amygdala E_Loc_ within the *Whole Brain* network.

**Conclusions:** Using a data-driven analysis approach, high-quality parcellation, and clinically diverse adolescent cohort, we found that symptoms within positive and negative valence system constructs differentially associated with resting-state network abnormalities: depression and anhedonia, as well as clinical status, involved greater influence and communication efficiency in prefrontal and limbic reward areas, whereas anxiety was linked to reduced influence/efficiency in amygdala and cortical regions involved in stimulus monitoring.

## Introduction

Adolescent depression represents a major public health concern and is associated with significant morbidity, including academic and occupational failure, social dysfunction, substance use, and, critically, suicide (1). Depression incidence rises dramatically during adolescence, with an estimated 15% of the population experiencing at least one major depressive episode between ages 15-24 (2). This increase has been attributed to deviations from normal maturational processes involving synaptic pruning, myelination, neurotransmission, and intrinsic functional circuits (3). In particular, the reward system undergoes extensive neurodevelopmental changes during adolescence, creating a window of vulnerability when abnormalities in reward processing can potentially arise (4).

While reward dysfunction is a core aspect of depression, it is a salient feature across other psychiatric conditions, yet is highly variable even within the same disorder. Our laboratory has documented that severity of anhedonia, a direct manifestation of reward deficits, is widely distributed even in adolescents with moderate-to-severe current depressive episodes: some report extremely low subjective hedonic experience, while others report little or no reward impairment (5). Similarly, we found that anhedonia and overall depression levels were associated with distinct striatum-based resting-state functional connectivity patterns in depressed youths (6). These results highlight the need to account for inter-individual variations in symptom severity during clinical research. Indeed, a persistent challenge in delineating the neural underpinnings of mental illness has been reliance on heterogeneous diagnostic categories: a given disorder may include patients who share few clinical features, while nominally distinct disorders often share several features (7). Mood and anxiety disorders are especially notorious for having high rates of comorbidity and symptom overlap. In response, our group and others have increasingly focused on specific symptoms, which represent narrowly defined clinical features with potentially distinct etiologies, rather than broad categorical diagnoses.

Separately, recent developments in neuroimaging methodology have significantly advanced the understanding of *in vivo* brain function and hold great promise to elucidate the biological underpinnings of mental illness. Much of this work has been spearheaded by the Human Connectome Project (HCP) (8), which released a landmark cortical parcellation in 2016 identifying 360 distinct areas based on multimodal measures of cortical thickness, myelination, resting-state functional connectivity, and task activation patterns in an extensively sampled cohort of healthy young adults (9). Recently, the Cole-Anticevic Brain-wide Network Partition (CAB-NP) has extended this parcellation scheme to the subcortex, identifying 358 further regions on the basis of resting-state network assignments and providing a detailed map of discrete functional areas across the entire brain (10). In addition to revealing fundamental aspects of neural organization, high-quality parcellations provide an invaluable framework for further data-driven research. Principled data reduction is especially crucial in graph theory, which models complex systems like the brain as collections of nodes (e.g. cortical areas) linked by edges (e.g. functional connectivity) (11). In addition to concisely representing resting-state networks, graph theory can reveal numerous subtle network features, including nodal measures of centrality (i.e. influence over other nodes) and efficiency (i.e. ease of communication with other nodes) that compliment and transcend the information provided by traditional connectivity analyses. Anomalous network properties have been reported across a wide range of psychiatric and neurological conditions (12). Unfortunately, widespread differences in methodology, overly simplistic network models, and functionally inaccurate node boundaries have contributed to inconsistent findings (13, 14), limiting the utility of graph theory for understanding mental illness to date.

In the current study, we sought to use high-quality parcellations and detailed network models to examine the neural correlates of depression, anhedonia, and anxiety, assessed quantitatively in psychotropic-medication-free adolescents with diverse psychiatric symptoms. As clinical symptomatology is salient across disorders and lies on a continuum within each disorder, our study was designed to capture the full range of symptom severity by recruiting a large, transdiagnostic sample that included adolescents with comorbid and subthreshold diagnoses as well as healthy controls. Using graph theory, we examined relationships between clinical symptomatology and resting-state network properties of centrality and efficiency within the functionally accurate CAB-NP network (10). As reward circuitry plays a central role in the emergence of psychiatric conditions during adolescence, we also repeated analyses within three functionally defined reward networks derived from the Reward Flanker Task (RFT) (15). We hypothesized that clinical status and symptom severity would be associated with resting-state network properties in regions related to reward processing.

## Methods and Materials

### Recruitment

Adolescents, ages 12-20, were recruited from the greater New York City area. The study was approved by the Institutional Review Board at Icahn School of Medicine at Mount Sinai (ISMMS). Prior to the study, procedures were explained to adolescents and legal guardians. Participants age 18+ provided written consent; those under 18 provided written assent and a guardian provided written consent.

### Inclusion and Exclusion Criteria

#### All Participants

Adolescents were excluded if they had any significant medical or neurological condition, estimated IQ<80, claustrophobia, any MRI contraindication, or a positive urine toxicology or pregnancy test.

#### Psychiatric Group

Clinical participants were psychotropic-medication-free for 30+ days, or 90+ days for long half-life medications (e.g. fluoxetine). Exclusionary diagnoses were pervasive developmental disorders, current psychosis, or a substance use disorder in the past year. All other psychiatric conditions were allowed, regardless of whether full diagnostic criteria were met.

#### Healthy Control Group

Control participants did not meet criteria for any current or past psychiatric diagnoses and were psychotropic-medication-naïve.

### Clinical Measures

#### Diagnostic Procedures

Clinical and sub-clinical DSM-IV-TR diagnoses were obtained using the Kiddie Schedule for Affective Disorders and Schizophrenia – Present and Lifetime Version (K-SADS-PL) (16). Interviews were administered to all adolescent participants, as well as a guardian if the participant was under 18. Evaluations were discussed between the interviewing clinician and Principal Investigator (VG), a board-certified child and adolescent psychiatrist, in order to enhance reliability.

#### Depression

Overall depression severity was assessed using the Beck Depression Inventory-II (BDI), a 21-item scale that assesses symptoms and features of depression over the previous two weeks. The BDI has high internal consistency in both clinical and non-clinical adolescent populations (17).

#### Anhedonia

Severity was assessed by the state-based Temporal Experience of Pleasure Scale (TEPS). This 18-item self-report separately quantifies anticipatory (TEPS-A) and consummatory (TEPS-C), as well as total (TEPS-T), anhedonia symptoms over the past week (18). Since the TEPS is reverse-scored (higher scores → lower anhedonia), analyses were performed using negative TEPS values (higher scores → higher anhedonia) for consistency with other scales.

#### Anxiety

Severity was examined using the Multidimensional Anxiety Scale for Children (MASC), a 39-item scale validated in both clinical and non-clinical populations (19).

### Imaging Data Acquisition

Data were acquired on a 3T Skyra MR system (Siemens, Germany) with 16/4-channel head/neck coil using protocols similar to the HCP Lifespan study (20). Sequences included: T1-weighted MPRAGE (TR=2400ms; TE=2.06ms; TI=1000ms; flip angle=8°; 224 frames, no gap; matrix=256×256; FOV=230×230mm^2^; 0.9mm isotropic), T2-weighted SPACE (TR=3200ms; TE=565ms; flip angle=120°; 224 frames, no gap; matrix=256×256; FOV=230×230mm^2^; 0.9mm isotropic), and resting-state gradient-recalled EPI (TR=1000ms; effective TE=31.4ms; flip angle=60°; 600 frames of 60 slices parallel to AC-PC, no gap; 5× multiband acceleration; anterior-to-posterior phase encoding; matrix=98×98; FOV=228×228mm^2^; 2.3mm isotropic; 10min). Matched single-band EPI and spin-echo fieldmaps were collected for registration and distortion correction purposes. Subjects were presented with a fixation cross and instructed to rest with their eyes open. Four RFT fMRI runs (15) were also acquired later in the session.

### Imaging Data Processing

Data were visually inspected before preprocessing with HCP Pipelines v3.4 (21). For anatomical data, preprocessing included gradient non-linearity correction, b0 distortion correction, AC-PC alignment, coregistration, brain extraction, bias-field correction, nonlinear transformation to MNI space, FreeSurfer segmentation, and cortical ribbon extraction. Functional data were corrected for gradient nonlinearity and EPI readout distortion, realigned, transformed to MNI space, intensity normalized, and mapped to the cortical ribbon. Structured noise components in concatenated resting-state and RFT fMRI runs were automatically identified via ICA-FIX classifier (22, 23), manually reviewed, and regressed out. Cortical surface data were robustly aligned across subjects based on a combination of functional and anatomical features using multimodal surface matching (MSMAll), developed and advocated by the HCP (24, 25). Resting-state data were further denoised by regressing out 24 movement parameters (6 affine + temporal derivatives + squares of each) and the five largest principle components associated with white matter and cerebrospinal fluid (CompCor), identified using Conn v17f (26). Finally, fMRI data were parcellated (see below) and bandpass filtered (0.1-0.01Hz). No spatial smoothing kernel was applied. All analyses were performed in 32k-CIFTI grayordinate space, which combines 2D left and right cortical surface representations with 3D subcortical data in MNI space.

Functional data were divided into and averaged within nodes using CAB-NP v1.0.5, which extends the HCP cortical parcellation (9) to include functionally similar subcortical parcels (10). As in previous work (27), we slightly modified the cortical parcellation by subdividing the somatomotor strip along somatotopic boundaries, yielding the final *Whole Brain* network (750 nodes). Additionally, we identified three reward-related networks (**Figure 1**) based on a separate analysis of RFT fMRI data collected in the same sample (manuscript in preparation), which builds on our previous RFT studies (15, 28). Briefly, these networks comprised the 10% of nodes most activated by *Reward Anticipation* (114 nodes), *Reward Attainment* (103 nodes), and *Reward Prediction Error* (117 nodes) RFT contrasts, as well as any corresponding contralateral nodes.

**Figure 1:**
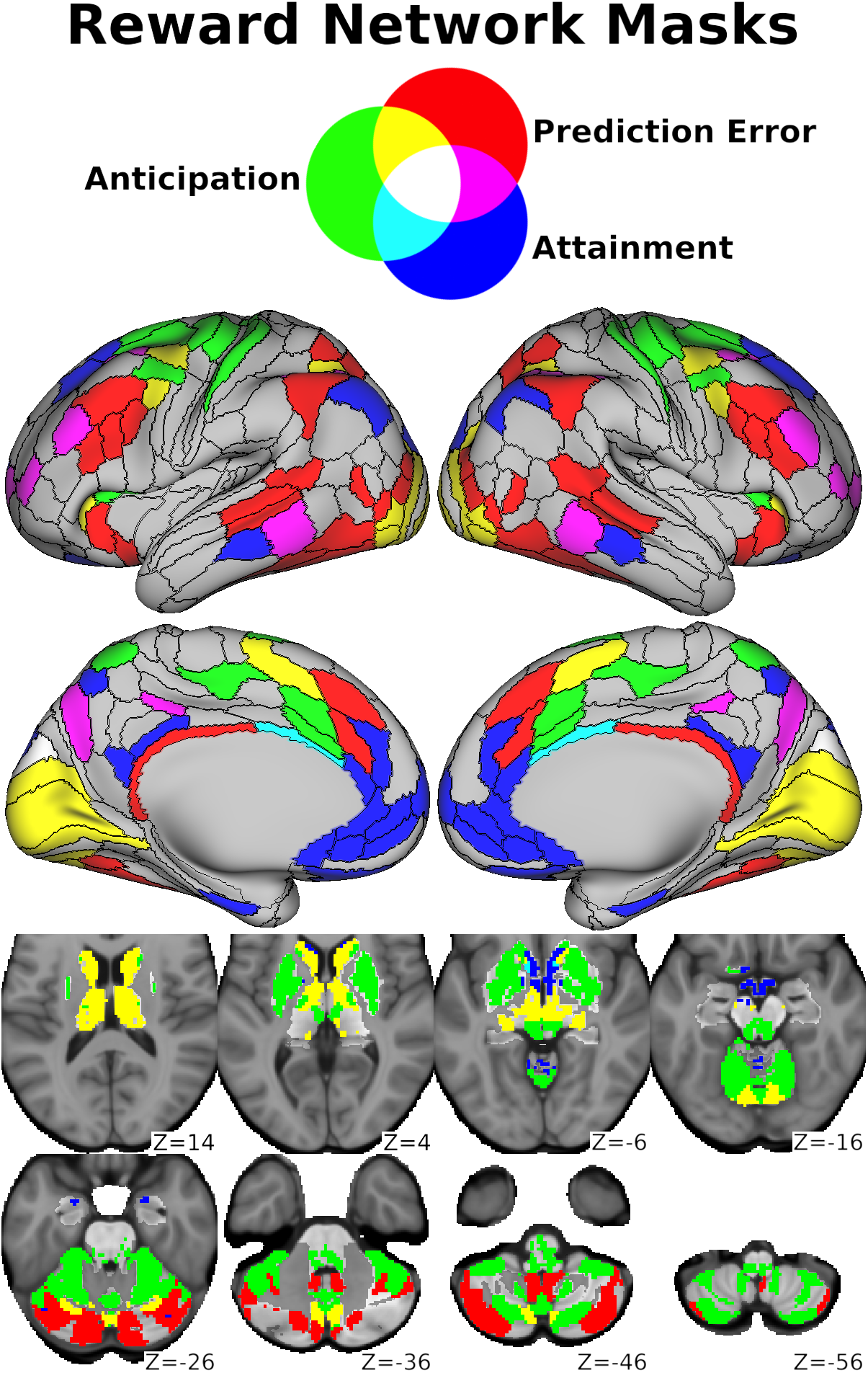
Nodes from the *Whole Brain* network corresponding to the *Reward Anticipation* (green), *Reward Attainment* (blue), and *Reward Prediction Error* (red) RFT networks. Nodes included in more than one network are indicated by additive color mixing, as shown at the top.

### Graph Theoretical Metrics

Subject-level association matrices were generated by cross-correlating node timeseries within the *Whole Brain* (750×750), *Reward Anticipation* (114×114), *Reward Attainment* (103×103), and *Reward Prediction Error* (117×117) networks in MATLAB v2017a and retaining all positive *r* values. Within each network, graph theoretical metrics were then estimated using weighted, undirected measures from Brain Connectivity Toolbox v2019-03-03 (11):

1. <u>Strength Centrality</u> (C_Str_): The sum of all edge weights (i.e. positive *r* values) at each node. C_Str_ is the weighted analogue of the binary Degree Centrality metric.
2. <u>Eigenvector Centrality</u> (C_Eig_): The eigenvector with the largest eigenvalue for each node. This measurement is self-referential, such that nodes with high C_Eig_ are those most closely associated with other high-C_Eig_ nodes.
3. <u>Local Efficiency</u> (E_Loc_): The inverse shortest path length (i.e. minimum number of edges, adjusted for edge weights) between each node and its neighborhood.

### Graph Theory Analysis

Group differences (clinical vs. control) in graph theoretical metrics were assessed using two-sample *t*-tests. Relationships between graph theoretical metrics and symptom scales (BDI, MASC, negative TEPS) were assessed using Pearson partial correlations in the full sample. All analyses controlled for participant age and sex. Statistical significance was determined using non-kwd-groupparametrickwd-grouppermutation tests (10,000 iterations), as implemented in FSL PALM v111alpha (29). Non-parametric tests provide better familywise error (FWE) control than their parametric equivalents (30) and are robust to skewed data distributions (31), as was the case for symptom scales in our study (skewness: BDI=1.32; MASC=0.58; TEPS-A=−0.94; TEPS-C=−0.75; TEPS-T=−1.00). Results were considered significant at the two-tailed *p*_*FWE*_<0.05 level. All results data are freely available through BALSA (32) at https://balsa.wustl.edu/study/show/x278x (data will be released upon peer-reviewed publication).

## Results

### Clinical Characteristics

The sample included 87 adolescents, of whom 68 had psychiatric symptoms and 19 were healthy controls. **Table 1** provides participant demographic and clinical characteristics. Relative to controls, adolescents with psychiatric symptoms had significantly higher BDI and MASC scores (*p*_*FWE*_<10^−3^). Groups did not differ significantly in age, sex, race, ethnicity, or TEPS scores (*p_FWE_*>0.1).

**Table 1:** Clinical and Demographic Information. Demographic and diagnostic information for the healthy control, clinical, and combined adolescent groups.

| **Table 1: Clinical and Demographic Information** |  |  |  |
| --- | --- | --- | --- |
| **Measure** | **Control (n=19)** | **Clinical (n=68)** | **All (N=87)** |
| Age (M ± SD) | 15.3 ± 2.5 | 15.1 ± 2.1 | 15.2 ± 2.2 |
| Sex | F=9, M=10 | F=44, M=24 | F=53, M=34 |
| Race <sup>a</sup> | Af=8, As=0, E=7, O=4 | Af=24, As=2, E=30, O=12 | Af=32, As=2, E=37, O=16 |
| Ethnicity <sup>b</sup> | H=5, N=14 | H=33, N=35 | H=38, N=49 |
| BDI (M ± SD) | 1.8 ± 2.1 (n=19) | 13.6 ± 11.4 (n=66) | 11.0 ± 11.2 (N=85) |
| MASC (M ± SD) | 27.0 ± 11.7 (n=17) | 44.7 ± 17.3 (n=66) | 41.1 ± 17.8 (N=83) |
| TEPS-A (M ± SD) | 49.4 ± 6.8 (n=18) | 45.0 ± 8.9 (n=52) | 46.1 ± 8.6 (N=70) |
| TEPS-C (M ± SD) | 35.0 ± 8.7 (n=18) | 33.2 ± 7.5 (n=52) | 33.6 ± 7.8 (N=70) |
| TEPS-T (M ± SD) | 84.4 ± 13.7 (n=18) | 78.1 ± 14.5 (n=52) | 79.7 ± 14.5 (N=70) |
| Mood Symptoms <sup>c</sup> | 0 | 49 | 49 |
| Anxiety Symptoms <sup>c</sup> | 0 | 43 | 43 |
| Behavioral Symptoms <sup>c</sup> | 0 | 28 | 28 |
| Other Symptoms <sup>c</sup> | 0 | 7 | 7 |
| <sup>a</sup> Af=African American, As=Asian American, E=European American, O=Other/Mixed Race<br><sup>b</sup> H=Hispanic, N=Non-Hispanic<br><sup>c</sup> Includes past and/or subthreshold symptoms |  |  |  |

### Group Differences

Adolescents with psychiatric symptoms had significantly higher C_Eig_ in both the left ventral striatum and right extrastriate visual cortex within the *Reward Attainment* network (**Figure 2**, **Table 2**). Group differences were non-significant for all other metrics and networks.

**Table 2:** Graph Theory Group Contrast and Symptom Correlation Results. Detailed results including parcellation labels, effect sizes, and two-tailed *p*_*FWE*_ values for each significant node.

| Table 2: Graph Theory Group Contrast and Symptom Correlation Results |  |  |  |  |  |
| --- | --- | --- | --- | --- | --- |
| Network | Metric | Location | CAB-NP Label (HCP Label)<sup>a</sup> | Effect Size<sup>b</sup> | p<sub>FWE</sub> |
| Clinical vs. Control Adolescents |  |  |  |  |  |
| Reward Attainment | C<sub>Eig</sub> | Right Ventral Striatum | Orbito-Affective-3 | 0.947 | 0.048 |
|  |  | Right Visual Cortex | Visual2_3 (V3) | 0.945 | 0.047 |
| Depression Severity |  |  |  |  |  |
| Whole Brain | C<sub>Str</sub> | Right sgACC | Default_34 (25) | 0.426 | 0.023 |
|  | E<sub>Loc</sub> | Right sgACC | Default_34 (25) | 0.392 | 0.045 |
| Anticipatory Anhedonia Severity |  |  |  |  |  |
| Whole Brain | C<sub>Str</sub> | Left vmPFC | Default_46 (10r) | 0.451<sup>c</sup> | 0.030 |
|  |  | Left Pulvinar Thalamus | Auditory-24 | 0.442<sup>c</sup> | 0.044 |
|  | E<sub>Loc</sub> | Left Pulvinar Thalamus | Auditory-24 | 0.469<sup>c</sup> | 0.013 |
|  |  | Left Central Amygdala | Dorsal Attention-1 | 0.432<sup>c</sup> | 0.048 |
|  |  | Right Lateral Amygdala | Posterior_Multimodal-2 | 0.434<sup>c</sup> | 0.044 |
| Total Anhedonia Severity |  |  |  |  |  |
| Whole Brain | E<sub>Loc</sub> | Left Pulvinar Thalamus | Auditory-24 | 0.431<sup>c</sup> | 0.050 |
| Anxiety Severity |  |  |  |  |  |
| Whole Brain | C<sub>Eig</sub> | Right dlPFC | Frontoparietal_12 (a9-46v) | 0.428 | 0.025 |
|  |  | Right Parietal Operculum | Cingulo-Opercular_16 (PFcm) | -0.413 | 0.044 |
|  | E<sub>Loc</sub> | Right Medial Amygdala | Somatomotor-3 | -0.390 | 0.050 |
| Reward Anticipation | C<sub>Str</sub> | Left Ventral Visual Stream | Visual2_40 (PIT) | -0.367 | 0.036 |
| Reward Prediction Error | C<sub>Str</sub> | Left Ventral Visual Stream | Visual2_36 (FFC) | -0.360 | 0.044 |
| <sup>a</sup> Labels per Cole-Anticevic Brain-wide Network Partition v1.0.5 (equivalent cortical labels per HCP S1200 release)<br><sup>b</sup> Cohen's *d* for group contrasts and Pearson's *r* for symptom correlations, adjusted for age and sex<br><sup>c</sup> Correlations reported with negative TEPS scores for consistency with other scales (see **Methods and Materials**) |  |  |  |  |  |

**Figure 2:**
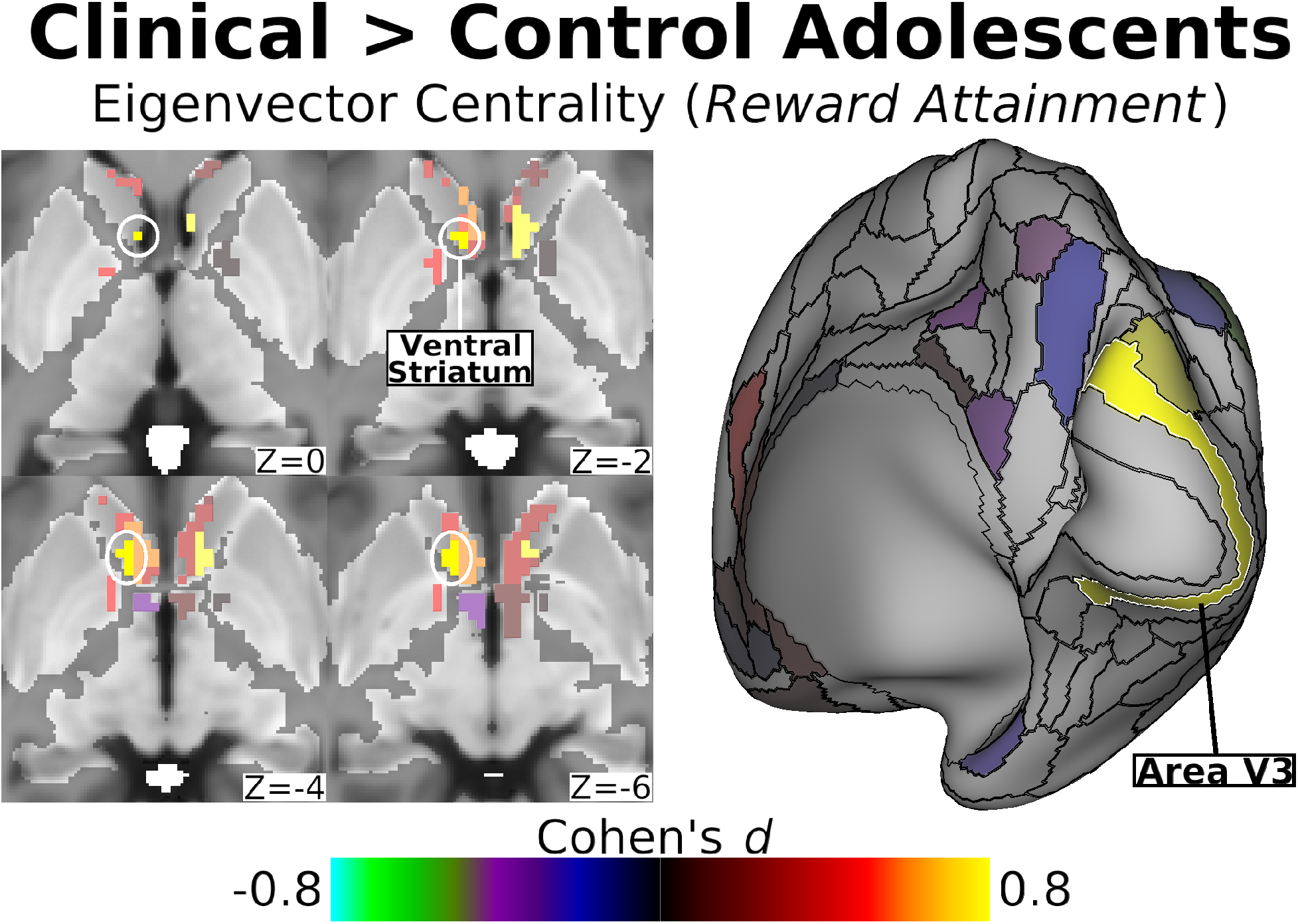
Within the *Reward Attainment* network, adolescents with psychiatric symptoms had significantly higher C_Eig_ in the left ventral striatum (left) and area V3 of the right extrastriate visual cortex (right) than healthy controls. Maps show effect size (Cohen’s *d*), adjusted for age and sex. Significant (two-tailed *p*_*FWE*_<0.05) nodes are indicated by white outlines and labels; non-significant nodes are displayed at 50% saturation.

### Depression Severity

Depression severity (BDI) was positively correlated with both C_Str_ and E_Loc_ in the right subgenual anterior cingulate cortex (sgACC) within the *Whole Brain* network (**Figure 3**, **Table 2**). No significant associations with depression were found for C_Eig_ or within RFT networks.

**Figure 3:**
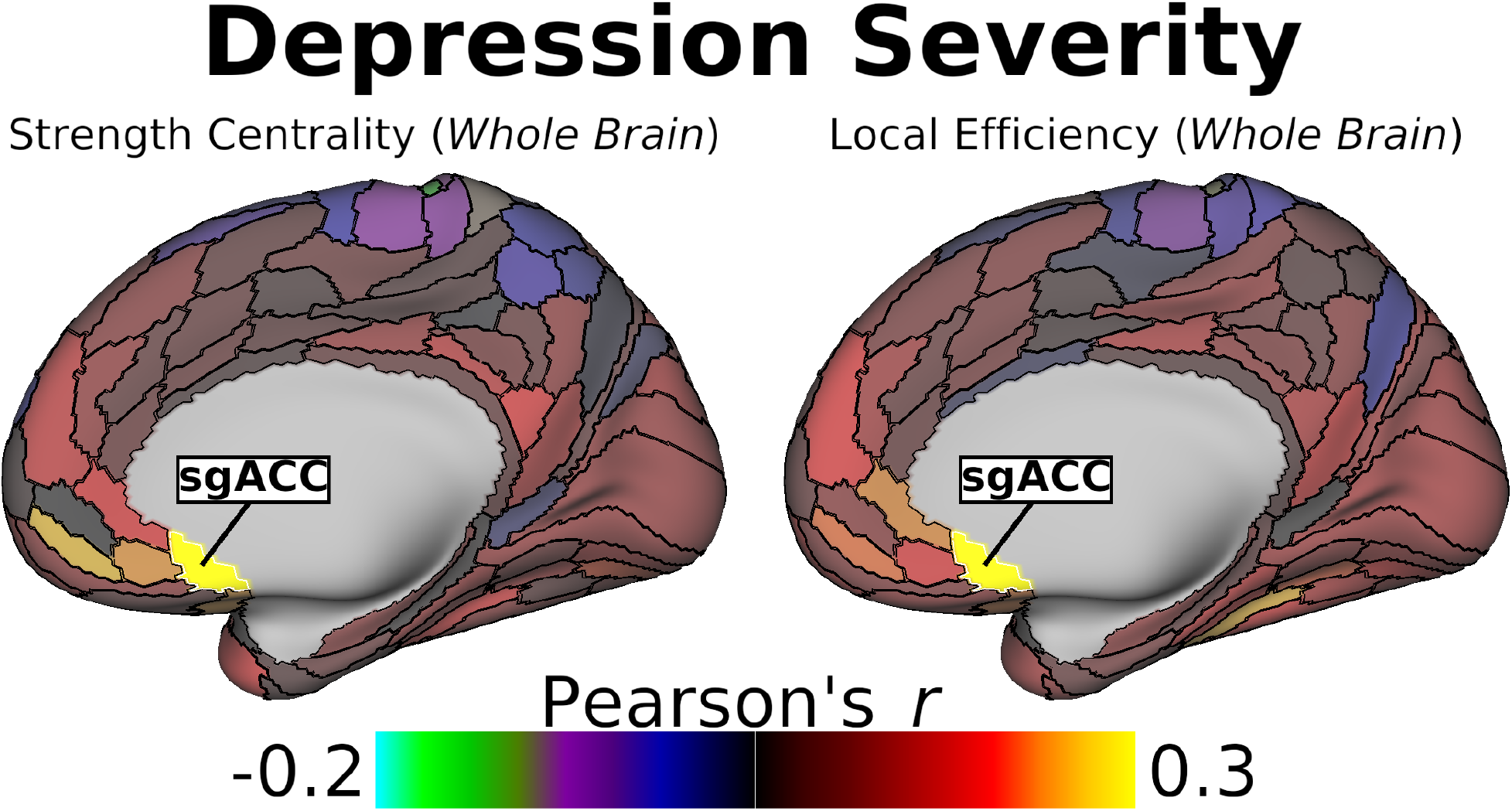
Within the *Whole Brain* network, overall depression severity was positively correlated with both C_Str_ (left) and E_Loc_ (right) in the right sgACC across all adolescents. Maps show effect size (Pearson’s *r*), adjusted for age and sex. Significant (two-tailed *p_FWE_*<0.05) nodes are indicated by white outlines and labels; non-significant nodes are displayed at 50% saturation.

### Anhedonia Severity

Within the *Whole Brain* network, anticipatory anhedonia severity (negative TEPS-A) was significantly correlated with C_Str_ in the left ventromedial prefrontal cortex (vmPFC) and left medial pulvinar thalamus, as well as with E_Loc_ in the right lateral amygdala, left central amygdala, and left medial pulvinar thalamus (**Figure 4**, **Table 2**). Total anhedonia severity (negative TEPS-T) was also significantly correlated with *Whole Brain* E_Loc_ in the left medial pulvinar thalamus (**Table 2**). No significant correlations with consummatory anhedonia severity (negative TEPS-C) or within RFT networks were detected.

**Figure 4:**
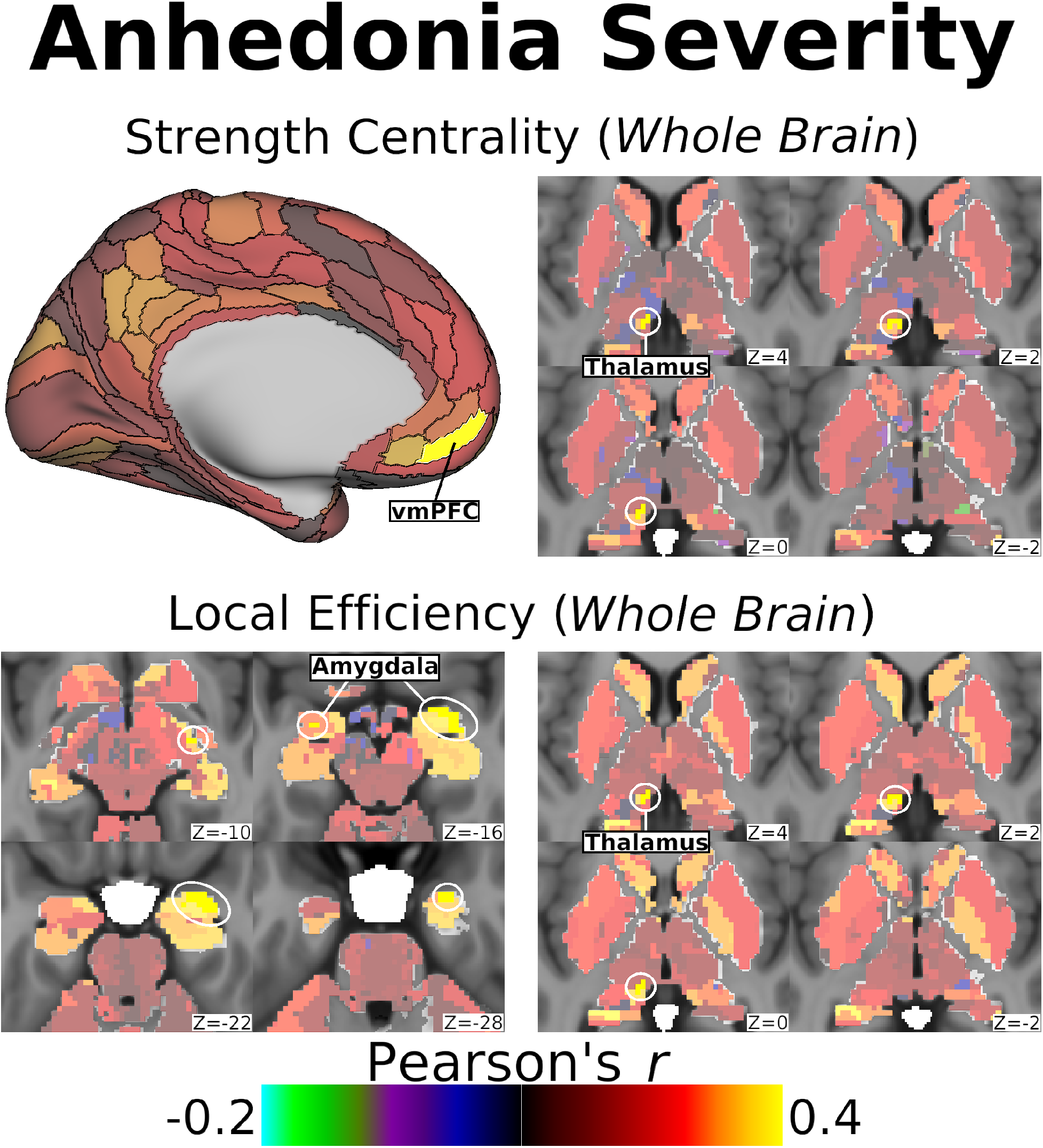
Within the *Whole Brain* network, anticipatory anhedonia was positively correlated with C_Str_ in the left vmPFC (top left) and left medial pulvinar thalamus (top right), as well as with E_Loc_ in the bilateral amygdala (bottom left) and left medial pulvinar thalamus (bottom right) across all adolescents. Maps show effect size (Pearson’s *r*), adjusted for age and sex. Significant (two-tailed *p_FWE_*<0.05) nodes are indicated by white outlines and labels; non-significant nodes are displayed at 50% saturation.

### Anxiety Severity

Within the *Whole Brain* network, anxiety severity (MASC) was positively correlated with C_Eig_ in the right dorsolateral prefrontal cortex (dlPFC), negatively correlated with C_Eig_ in the right parietal operculum, and negatively correlated with E_Loc_ in the right medial amygdala (**Figure 5**, **Table 2**). Additionally, anxiety was negatively correlated with C_Str_ in parts of the left ventral visual stream, specifically the PIT Complex within the *Reward Anticipation* network and the adjacent FF complex within the *Reward Prediction Error* network. No significant anxiety correlations were found within the *Reward Attainment* network.

**Figure 5:**
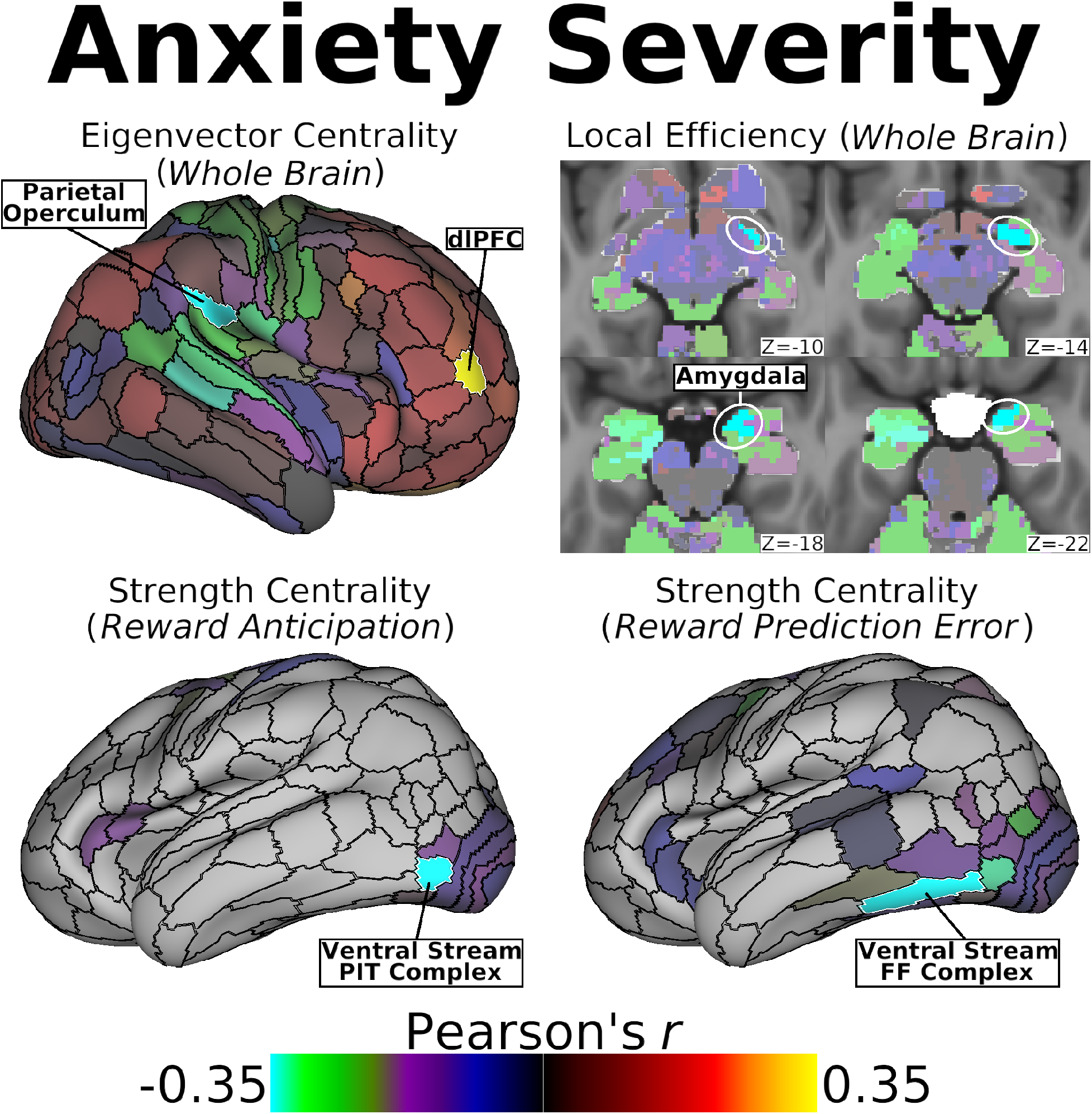
Within the *Whole Brain* network, anxiety severity correlated positively with right dlPFC C_Eig_ and negatively with right parietal operculum C_Eig_ (top left), and also correlated negatively with E_Loc_ in the right medial amygdala (top right) across all adolescents. Additionally, anxiety was negatively correlated with C_Str_ in two left ventral visual stream nodes: the PIT Complex within the *Reward Anticipation* network (bottom left) and the adjacent FF Complex within the *Reward Prediction Error* network (bottom right). Maps show effect size (Pearson’s *r*), adjusted for age and sex. Significant (two-tailed *p_FWE_*<0.05) nodes are indicated by white outlines and labels; non-significant nodes are displayed at 50% saturation.

## Discussion

The present study capitalized on recent advances in neuroimaging methodology to examine resting-state network properties in the context of adolescent mental illness. Our approach included high-quality multiband fMRI sequences to achieve excellent spatial (2.3mm isotropic) and temporal (1s) resolution, HCP-style preprocessing including highly accurate MSMAll surface alignment, and a large sample of psychotropic-medication-free adolescents with diverse clinical symptomatology. A key element of our study was the CAB-NP parcellation, which enabled us to model networks using functionally discrete nodes across the entire cortex and subcortex. To further preserve neurobiological detail, we derived graph theoretical metrics of centrality (C_Str_, C_Eig_) and efficiency (E_Loc_) using weighted association matrices, rather than the simpler binary approach where association matrices are arbitrarily thresholded and all surviving correlations are treated as equivalent. In addition to *Whole Brain* analyses, we also examined graph theoretical metrics within specific *Reward Anticipation*, *Reward Attainment*, and *Reward Prediction Error* networks, which we defined empirically using task fMRI data collected in the same subjects. Importantly, these analyses within smaller RFT networks did not simply reduce multiple comparison penalties, as in small-volume correction (33), but directly altered the calculation of graph theoretical metrics by restricting the underlying association matrix to nodes involved in the corresponding reward process.

As hypothesized, findings from both the group comparison and symptom correlation analyses implicated key reward-related areas, supporting the notion that alterations in reward circuitry during adolescent brain development play an important role in the emergence of psychiatric disorders. Specifically, adolescents with clinical symptoms vs. controls had significantly higher C_Eig_ in the ventral striatum within the *Reward Attainment* network. Across all adolescents, moreover, higher depression severity was associated with increased *Whole Brain* C_Str_ and E_Loc_ in the sgACC, while higher anhedonia severity was associated with the increased *Whole Brain* C_Str_ in the vmPFC. Taken together, these findings suggest that elevated tonic (i.e. resting-state) communication with reward areas may be related to the initial development of positive valence system (PVS) deficits. By contrast, the negative valence system (NVS) construct of anxiety was mainly associated with reduced tonic communication with areas related to salience and threat monitoring.

Our group-level results highlight the importance of studying specific reward sub-systems, especially for a heterogeneous cohort: we found that adolescents with mood, anxiety, and/or behavioral disorder symptoms were only distinguishable from their healthy counterparts based on network topology within the *Reward Attainment* network. Within this network, clinical adolescents had elevated C_Eig_ in the left ventral striatum, specifically the ventromedial caudate bordering the nucleus accumbens (NAc). The ventral striatum plays a highly conserved role in primary reward processing, receiving dopaminergic inputs from the ventral tegmental area in response to pleasant stimuli via the mesolimbic reward pathway (34, 35). Similar to our results, a previous resting-state fMRI study in children aged 6-12 found that, within a network consisting of 12 reward-related nodes, only left ventral striatum C_Str_ significantly predicted the emergence of depression and was associated at the trend level with development of ADHD and anxiety at 3-year follow-up (36). Our finding of significant differences in left ventral striatal C_Eig_ (i.e. association with influential nodes), rather than C_Str_ (i.e. association with all nodes), may be due to the much larger number of nodes in our *Reward Attainment* network or differences in C_Str_ calculation. In addition to the ventral striatum, we also detected significantly higher C_Eig_ in area V3 of the right visual cortex in the clinical cohort. Although the extrastriate visual cortex is unlikely to play a direct role in reward processing, reward responses are often contingent on sensory inputs, and area V3 was notably the only node to appear in all three reward networks (white in **Figure 1**). The lack of significant group differences in other networks may have been related to the heterogeneity of the clinical group.

Supporting this conclusion, clinical symptoms were primarily associated with the resting-state properties of nodes within the *Whole Brain* network. Specifically, overall depression severity was positively correlated with both C_Str_ and E_Loc_ in the right sgACC, an area that is heavily implicated in depression. Increased sgACC activity is frequently reported in neuroimaging studies of depressed adults (37) and adolescents (38), while sgACC activity decreases following many types of depression treatment, including traditional antidepressants, ketamine, and deep brain stimulation (39–41). Resting-state fMRI studies have indicated that depression severity is associated with sgACC connectivity changes in adults with clinical (42) and subclinical (43) depression as well as depressed adolescents (44, 45), which may be related to early-life changes in white mater microstructure (46, 47). Our findings add to this body of evidence, showing that higher overall sgACC connectivity (C_Str_) and shorter connectivity paths to the sgACC (E_Loc_) are associated with increased depression severity across a large cohort of clinically diverse adolescents.

In addition to overall depression levels, our analyses also revealed distinct correlations between *Whole Brain* network properties and anhedonia severity. Of note, these findings were driven by anticipatory anhedonia, which involves undervaluation of expected rewards and is associated with motivational deficits, rather than consummatory anhedonia, which reflects diminished experience of pleasure once rewards are obtained. Together with our group contrast results, which only detected significant differences within the *Reward Attainment* network associated with reward consummation, this finding highlights the importance of considering discrete phases of reward processing, even at rest. We found that anticipatory anhedonia severity correlated with overall vmPFC connectivity (C_Str_) and with shorter connectivity paths to the mid-lateral amygdala (E_Loc_). The vmPFC receives extensive reward-related inputs from the dopaminergic midbrain and ventral striatum via the mesocorticolimbic pathway (48) and is responsive to many types of primary rewarding stimuli (49). Previous studies have reported a link between anticipatory anhedonia and reduced vmPFC activation to rewarding stimuli regardless of diagnosis (50), and reduced vmPFC connectivity with subcortical reward structures is associated with anhedonia as well as inflammation in depression (51). Moreover, a transcranial magnetic stimulation treatment study in adults with depression found that non-responders relative to responders had both higher anhedonia scores and higher betweenness centrality (a related graph theoretical measure) only in the vmPFC (52). Reduced amygdala resting-state functional connectivity with the vmPFC has also been repeatedly reported in adolescents with depression relative to controls (53, 54), whereas the opposite pattern was observed in a large meta-analysis of depressed vs. healthy adults (55), suggesting a potential effect of illness chronicity or treatment on these regions. Although the amygdala has been a frequent target of fMRI research, a major strength of our approach was the ability to discriminate between functionally distinct subregions of this and other heterogeneous subcortical structures. In particular, the basolateral nucleus of the amygdala plays a well-characterized role in motivation through glutamatergic projections to the ventral striatum, which converge on many of the same reward-encoding cells in the nucleus accumbens that receive mesolimbic dopamine inputs (56). Consistent with this, our analyses specifically identified a link between the lateral amygdala and anticipatory anhedonia in adolescents.

Our anxiety analyses, meanwhile, revealed the opposite relationship with E_Loc_ in the medial amygdala, in line with extensive literature tying the amygdala to fear, anxiety, and related NVS constructs (57). Others have previously reported amygdala functional connectivity was reduced with the orbitofrontal cortex/vmPFC in adults with social anxiety disorder (58) and with the anterior cingulate and insula in adults with generalized anxiety disorder (59). In adults with depression, reduced amygdala connectivity with the dorsomedial PFC, mid-/posterior cingulate, and lateral temporal areas was predictive of comorbid anxiety (60), and amygdala-vmPFC connectivity negatively correlated with anxiety levels (61). Possibly related to its unique position as an NVS, rather than PVS, construct, anxiety was the only symptom to have predominantly negative associations with graph theoretical metrics in our study. In addition to the amygdala, anxiety was anticorrelated with *Whole Brain* C_Eig_ in a parietal operculum node associated with the salience (a.k.a. cingulo-opercular) network, a group of regions involved in identifying and directing attention towards important stimuli. Due to its role in monitoring imminent threats as well as potential rewards, the salience network has been frequently implicated in both NVS and PVS dysfunction (62). The association between anxiety and brain regions involved in stimulus monitoring may also account for our findings in the ventral visual stream. Per the two-stream model, the ventrolateral occipital and temporal cortices form a “ventral stream” preferentially involved in determining the identity and salient characteristics of objects, whereas dorsolateral occipital and parietal areas form a “dorsal stream” primarily aimed at locating objects in space (63). In our study, adolescents with higher anxiety had lower overall connectivity (C_Str_) with ventral stream nodes within both the *Reward Anticipation* network, which is engaged during periods expectation when a reward has yet to be received, and the *Reward Prediction Error* network, which is differentially responsive to uncertain vs. certain reward attainment (15). As such, our findings indicate that brain regions important to externally oriented tasks of salience monitoring and reward discrimination have reduced tonic influence in adolescents with high anxiety levels.

As always, several caveats should be noted for this study. Foremost, although we recruited a relatively large cohort of 87 adolescents, sampling was more limited within major clinical categories of mood symptoms (n=49), anxiety symptoms (n=43), behavioral symptoms (n=28), and especially healthy controls (n=19). This study design was intended to capture the full range of clinical symptomatology by including subjects with significant comorbidity and subthreshold symptoms. As such, analyses focused primarily on associations with symptom severity in the full cohort; additional research is needed to determine how resting-state network properties differ between specific diagnostic groups and healthy adolescents. Second, although symptom severity is a more specific indicator of underlying PVS and NVS abnormalities than categorical diagnosis (7), clinical symptoms are also heterogeneous to some extent. We were able to address this directly for anhedonia by separately analyzing anticipatory and consummatory TEPS subscales; our future work will employ more granular assessments of depression and anxiety symptoms to allow comparable analyses. Finally, although we used the best whole-brain parcellation currently available, there has been limited validation of the CAB-NP due to its recent release. However, all cortical boundaries were taken directly from the multimodal surface parcellation meticulously derived by the HCP (9), which has been found to outperform other contemporary atlases and is widely considered a gold standard of human brain segmentation (64, 65). Subcortical parcels in the CAB-NP were then determined using a consensus partitioning approach based on data from over 300 HCP subjects divided into independent discovery and validation sets to ensure reproducibility and reliability (10).

In conclusion, our study prioritized high-quality clinical and neuroimaging measures, recruiting a large cohort of psychotropic-medication-free adolescents to examine the full range of illness severity using sophisticated fMRI acquisition and analysis techniques. We found that PVS constructs of depression and anhedonia severity as well as clinical status were associated with increased tonic communication with key reward-related nodes in the medial PFC and ventral striatum. Conversely, the NVS construct of anxiety was linked to reduced communication metrics in regions important to threat detection and stimulus monitoring. These results showcase the power of carefully constructed network models and data-driven analyses to detect specific functional anomalies underlying emergent clinical symptoms. Identifying and characterizing these aberrant neurodevelopmental processes is crucial for understanding and ultimately stopping the course of mental illness.

## Acknowledgements

This study was supported by National Institute of Mental Health (NIMH) grants R01MH101479 and R01MH095807 to VG. This work was also supported in part through the computational resources and staff expertise provided by ISMMS Scientific Computing, with additional resource support provided by the ISMMS Brain Imaging Center. We would like to thank Yael Jacob, PhD, for her advice regarding graph theory analysis.

## Disclosures

All authors declare no conflicts of interest.

## Notes

https://balsa.wustl.edu/study/show/x278x

